# MorphOT: Transport-based interpolation between EM maps with UCSF ChimeraX

**DOI:** 10.1101/2020.09.08.286302

**Authors:** Arthur Ecoffet, Frédéric Poitevin, Khanh Dao Duc

## Abstract

**Motivation:** Cryogenic Electron-Microscopy offers the unique potential to capture conformational heterogeneity, by solving multiple 3D classes that co-exist within a single cryo-EM image dataset. To investigate the extent and implications of such heterogeneity, we propose to use an optimal-transport based metric to interpolate barycenters between EM maps and produce morphing trajectories. While standard linear interpolation mostly fails to produce realistic transitions, our method yields continuous trajectories that displace densities to morph one map into the other, instead of blending them.

**Implementation:** Our method is implemented as a plug-in for *ChimeraX* called *MorphOT*, which allows the use of both CPU or GPU resources. The code is publicly available on GitHub (https://github.com/kdd-ubc/MorphOT.git), with documentation containing tutorial and datasets.

## Background

Cryogenic electron microscopy (cryo-EM) has yielded over the past decade a revolution in structural biology and the study of conformational heterogeneity [1]. To visualize the possible motion between conformational states solved by cryo-EM, a first approach is to generate continuous deformation of density maps with morphing-based methods. In *ChimeraX* –a standard visualization tool for EM maps [2]–, morphing is implemented with the *Volume Morph* package, which applies a linear interpolation to produce a trajectory joining two EM density maps. However, this linear interpolation can result in trajectories that unrealistically merge densities, instead of displaying continuous motion. We propose an alternate approach, based on the theory of optimal transport (OT) [3], that computes “displacement” interpolants [4] of EM maps. Implemented as a plug-in for *ChimeraX* called *MorphOT,* our bioinformatic tool mitigates the issues inherent to linear morphing, and can therefore help studying and visualizing transitions between conformational states.

## Description

To compute a continuous trajectory between two EM maps *V*_0_ and *V*_1_ (encoded as voxels in a 3D grid), we propose to *interpolate* between them by computing their weighted barycenters *V_t_* (0 < *t* < 1). In *ChimeraX*, the command *volume morph* computes *linear interpolants* 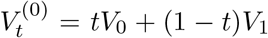. Instead of computing these interpolants, we perform with *MorphOT* a “displacement interpolation” [4,5], based on the so-called 2-Wasserstein distance 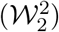. This distance can be seen as the minimal amount of work needed to move one mass density to another, with respect to a given cost function. For the two input EM maps *V*_0_ and *V*_1_ (seen as continuous or discretized distributions on 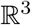), the *displacement interpolant* can then be written as [5]

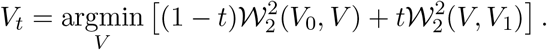

As this interpolation was first introduced to solve a partial advection problem [5], it typically characterizes some continuous motion from one density to the other. As a result, displacement interpolants tend to be more physically consistent, in contrast with blended solutions produced by linear interpolation [4].

More recently, Solomon *et al*. used entropic regularization to approximate this metric, leading to simple numerical schemes that efficiently evaluate these displacement interpolants, [3]. *MorphOT* follows the same approach to compute barycenters of EM maps, by combining these optimization algorithms with efficient implementations of Gaussian convolutions [6] (for more details, we refer to the user manual available on GitHub). As a plug-in for *ChimeraX, MorphOT* has been developped in Python 3.7, with non-ChimeraX pure Python source code also available on GitHub. The plug-in can run both on CPU or GPU, allowing large high resolution maps to be treated as well. Although a Python library for optimal transport already exists [7], it does not include convolutional Wasserstein distance computation, and is limited to 2D distributions. In particular, there has so far not been any implementation of the method proposed by Solomon *et al*. that is suitable to the typical format and size of EM maps. To our knowledge, our tool is thus the first implementation in Python able to deal with 3-dimensional grids of arbitrary size.

We illustrate in Figure 1 the results of *MorphOT* with two EMDB entries that correspond to two conformations of Mm-cpn, an archaeal group II chaperonin [8], showing back-to-back rings which can either be in an open or closed state. After pre-processing the maps by smoothing and denoising, we compared in Figure 1B the interpolation realized by *MorphOT* and ChimeraX’s *volume morph*, using three intermediate states (corresponding to interpolants *V*_1/4_, *V*_1/2_, *V*_3/4_). *Volume morph* superimposes the two inputs, with varying intensities for each interpolant. As a result, the rings from the open state already appear in *V*_1/4_, with *teleportation of mass* occurring between closed and open conformations. In contrast, *MorphOT* produces gradual openings of each closed branch from the rings. In agreement with the properties of the transport-based metric, we observe a *displacement* of mass from one location to the other.

**Figure 1:**
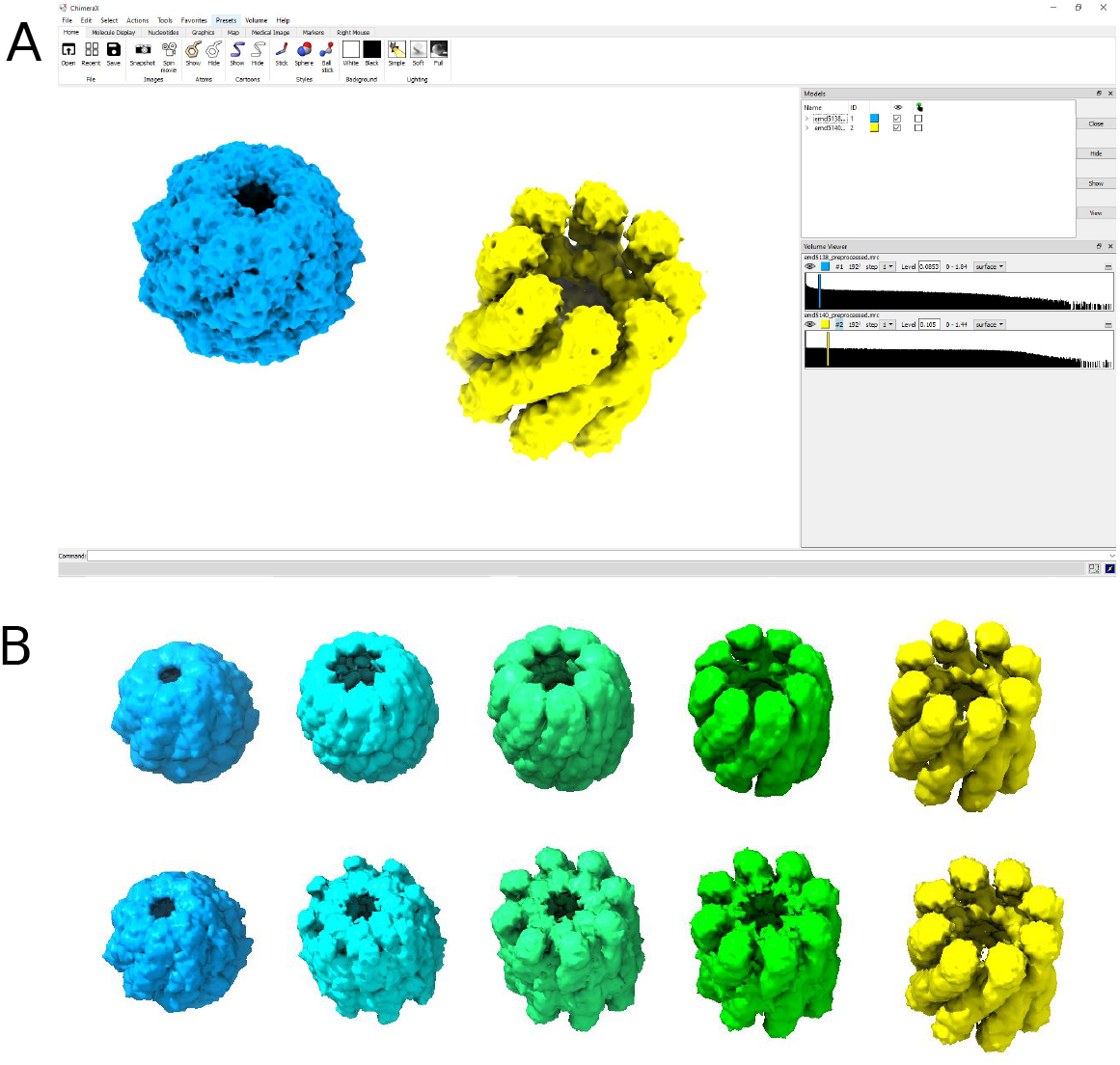
Interface of *MorphOT* and comparison with linear interpolation. **A:** ChimeraX’s interface showing on the left a closed state of lidless Mm-cpn, and on the right its open state (EMDB entries 5137/5139). **B:** Interpolants *V*_0_, *V*_1/4_, *V*_1/2_, *V*_3/4_, *V*_1_ obtained using *MorphOT* (Top) and *volume morph* (Bottom).

More generally, *MorphOT* takes two *ChimeraX* volumes (.mrc,.map,…) as main inputs, and allows to compute and display either the movie of the interpolating trajectory, or simply one barycenter. Barycenters can also be exported for further analysis. To mitigate the computational cost inherent with larger maps, we provide a faster approximation (*semiMorphOT*), as a mixed solution that computes a fraction of OT barycenters and linearly interpolates between them. Our user manual, available on the software GitHub page, contains a full detailed documentation of the algorithm and interface commands, with a tutorial that allows to reproduce the results shown here.

## Conclusion

MorphOT provides a new approach to visualize continuous trajectories between two EM maps. While this morphing tool generates physically plausible transitions, it is important to stress that further steps would be needed to validate these trajectories as ground truth. In this regard, it would be interesting to use MorphOT in complement to other approaches aiming to model continuous heterogeneity in cryo-EM data [9], and improve its accuracy by customizing the OT cost function using information from experimental data. We are currently pursuing these directions.

## Acknowledgements

KDD’s research is supported by NSERC DGECR-2020-00034 and NFRFE-2019- 00486 grants. We thank Dr. Simcha Srebnik for early discussion.

## Notes

https://github.com/kdd-ubc/MorphOT

### Competing Interest Statement

The authors have declared no competing interest.

